# Vocal sequences of diverse parrots suggest an important role for note repetition

**DOI:** 10.1101/2025.10.24.684295

**Authors:** Arjun Mandyam Dhati, Nakul Wewhare, Neeraj S. Devasthale, Saranya Sundar, Nihar Parulekar, Anand Krishnan

## Abstract

Sequencing and syntax in complex animal vocal signals have been studied using several mathematical analyses. Commonly, sequences of vocalizations are assumed to follow a first-order Markov process, where each state depends on the state immediately before it. However, more recent computational analyses challenge this assumption, suggesting alternative processes may be important in vocal sequences. Open-ended vocal learners such as parrots possess complex, variable vocal sequences that may be dynamically modified throughout their lives and contain information about group membership and identity. Although this makes them important systems in which to identify general patterns within vocal sequences, parrot vocalizations remain generally understudied compared to passerine birds. Here, we examined vocal sequence structure in six species of parrots (Psittaculidae). We fit various metrics of sequence structure to those generated by multiple simulated processes, including Markov, random, hidden Markov and renewal processes. Two metrics exhibited the best fit to a renewal process (where a note repeats a certain number of times before transitioning to a new note), and two others to Markov processes. Importantly, all analyses were broadly concordant across species, and multiple metrics indicated an elevated probability of note repetition. Reconciling these results, we suggest a general vocal mechanism across the highly variable vocal sequences of parrots, where both note repetition and Markov processes are important. Note repetition patterns could help communicate individual or group identity, as well as social and behavioral context. Our study thus extends a simulation-based approach using diverse metrics to comparatively examine the complex, variable vocal sequences of these open-ended vocal learners.

## Introduction

All forms of animal communication involve the transfer of information, and acoustic communication is no exception (1). Many animals sequence individual acoustic units into complex patterns, which may be modified to communicate information about behavioral state, context, individual and group identity (2–11). The study of complex communication signals has the potential to inform our understanding of vocal learning, speech recognition and cognition (7,12–14). Across diverse animal taxa, vocal sequences consist of individual units (frequently notes or syllables) that are emitted with some form of temporal ordering (7). The nature of this ordering has been examined using a number of mathematical methods, most commonly assuming a first-order Markov process using contingency tables (15,16). Here, the probability of finding any note (or state) is entirely determined by the state that immediately precedes it. However, recent research has suggested that many animal sequences do not follow a standard first-order Markov process (17–19). This has led to the proposal of alternative mechanisms structuring vocal sequences and varied analytical tools to examine these processes. First-order Markov processes rely on single-step transition probabilities, capturing first-order dependencies, whereas note co-occurrence (or collocation) metrics may be able to capture higher-order dependencies, and thus be better suited to analyzing complex sequences (9,20–22).

Birds produce some of the best-studied vocal sequences, typically in the form of songs for courtship and territorial advertisement (23). Many bird species learn song sequences at an early age, crystallizing them into a fixed pattern, whereas others learn and continuously modify their vocalizations throughout their lives (open-ended learning) (14,24,25). Learning processes result in syntactic or syllable-level changes in songs, which manifest as individual signatures or geographic dialects (5,13,20,26–33). Open-ended learners continuously modify songs throughout their lives, often at very short time scales (29,34,35). However, the often-variable vocal sequences in these taxa as well as their syntactic organization remain poorly studied. The order Psittaciformes, or parrots, are social, open-ended vocal learners with complex vocalizations and vocal sequences (3,5,6,14,24,29– 31,33–55). The acoustic structure of contact calls and the length of note repeats in vocal sequences encode both individual identity and the identity of the social group, and converge by social learning when groups merge or by individual matching (5,29–31,33,34,36,47–49,52,55). Vocal sequences are also variable and modified dynamically by behavioral events such as courtship displays or affiliative behavior (43). Despite the multiple sources of variability and complexity in parrot vocal sequences, we do not yet possess information on whether parrots exhibit broad, generalizable rules of vocal sequence ordering.

Here, we examined vocal sequence structure in six species of parrots from the family Psittaculidae (56), using multiple analytical methods and metrics. We recorded vocal sequences from captive birds held in a zoological collection. Employing a simulation-based framework (7,17), we tested whether rules governing parrot vocal sequences are generalizable. To do this, we generated simulated sequences assuming various processes, including a random sequence generator, a first-order Markov process, a renewal process and a hidden Markov process, and compared these to the observed sequences. We calculated multiple complementary metrics across the different processes and aimed for agreement across them. In doing so, we modified the rationale employed by Kershenbaum and Garland (2015) (57), aiming to examine whether multiple metrics could provide consensus on the process generating vocal sequences in parrots.

## Materials and Methods

### Recordings

We recorded vocal sequences from six different parrot species of the family Psittaculidae, native to diverse geographic regions (56): *Psittacula eupatria, Psittacula krameri, Psittacula cyanocephala, Agapornis nigrigenis, Agapornis roseicollis* and *Melopsittacus undulatus*. All species except *M. undulatus* were recorded at the Kamla Nehru Prani Sangrahalaya, Indore, Madhya Pradesh, India, between 0800 and 1200 hours. *P. krameri* and *P. eupatria* were housed in separate 30x15x15 foot aviaries, whereas the two *Agapornis* species were housed together in a single aviary of similar dimensions. *P. cyanocephala* was housed in a walk-in aviary with multiple other parrot species. Logistical and housing constraints prevented us from recording individual birds in separate cages, and so we recorded species within these aviaries, using video footage taken from a Nikon D3300 DSLR Camera (Nikon Corporation, Tokyo, Japan) equipped with 18-55mm and 70-300mm lenses to ensure that a single individual was emitting all the notes in a sequence. We used an ME66 directional microphone (Sennheiser Inc., Wedemark, Germany) attached to an H6 handy recorder (Zoom Corporation, Tokyo, Japan) to record vocalizations. The directionality of this microphone ensured a good signal:noise ratio in crowded aviary conditions. The number of individuals of each species held in the cages was as follows: *P. eupatria*: 33, *P. krameri*: 43, *P. cyanocephala*: 8, *A. roseicollis*: 18, *A*.*nigrigenis*: 14. The vocalizations of *M. undulatus* (5 males) were recorded as part of a previous study (43), and were added to this dataset.

### Annotating sequences

We used Raven Pro 1.6 (Cornell Laboratory of Ornithology, Ithaca, NY, USA) to label notes for all species. In total, our dataset contained 1859 notes from *P. eupatria* (268 sequences), 5493 notes from *P. krameri* (414 sequences), 526 notes from *P. cyanocephala* (48 sequences), 616 notes from *A. nigrigenis* (57 sequences), 606 notes from *A. roseicollis* (70 sequences), and 18171 notes from *M. undulatus* (411 sequences). We classified notes into categories based on spectral features (Supplementary Table 1) (5,20,43,58–60). All note classifications and annotations were performed by multiple authors, and the inter-observer agreement across species was consistent: *P. eupatria* - 95.35%; *P. krameri* - 82.74%; *P. cyanocephala* - 95.23%; *A. nigrigenis* - 88.2%; *A. roseicollis* - 89.80%; the classification scheme for *M. undulatus* was verified in previous studies (5,43).

We defined any group of vocalizations containing more than 3 notes as a vocal sequence. In order to account for the effects of variable gaps between notes, we defined a ‘silent period’ ‘q’, following a method from previously published studies (5,9,43). For each species, this was three times the maximum mean note duration across all note types: *P. eupatria*: 1s, *P. krameri*: 0.942s, *P. cyanocephala*: 0.777s, *A. nigrigenis*: 0.585s, *A. roseicollis*: 0.591s, and *M. undulatus*: 0.5s. Any case where there were two consecutive silent periods was considered a break in the sequence, and subsequent vocalizations were considered a separate sequence.

### Analysis

To examine which processes best explained the observed structure of parrot vocal sequences, we first quantified associations between notes using note co-occurrence matrices (9,21,22). Briefly, wedefined _d_C_ij_ as the probability that note type j occurs within d−1 notes of note type i. For all sequences in our dataset, we computed _d_C_ij_ values. Next, we generated randomized sequences where theprobability of occurrence of each note was given by its relative frequency of occurrence in the dataset for each species. The length of each sequence in this randomized dataset was randomly sampled from the lengths of recorded sequences, to ensure that the simulated sequences better approximated the observed data, and without artifacts resulting from unrealistically long sequences. From the randomized dataset, we computed _d_C_ij_ values, giving us the co-occurrence probability for this dataset. We then computed the odds-ratios across all species, defined as the ratio of the probability of two notes co-occurring in the recorded sequences to that of two notes co-occurring in the randomized dataset (9). Following this, we log-transformed the odds-ratios to have a uniform scale for comparing those ratios less than 1 and those greater than 1. A pseudocount of 0.01 was added to all values to avoid problems associated with logarithmic transformation for values equal to zero (9). We performed this analysis with d = 5; other studies have shown that the choice of d value does not significantly alter patterns within the matrix (5,9,20).

To understand the underlying generative process influencing sequence structure, we simulated sequences according to four different mathematical processes for each species (1000 simulated sequences per process per species, with the length of each sequence randomly sampled from the lengths of sequences in our dataset) and used diverse metrics to identify the best fit to the observed sequences. These processes were:

1. The sequences were random, with the probability of each note occurring at any position in the sequence being the same as its overall frequency of occurrence in the observed dataset.
2. The sequences were generated as first-order Markov chains. Here, each note emitted in a sequence depends solely on the note preceding it. We generated a transition probability matrix (denoted by M_0_) for each species, where the probability of transitioning from notes i to j, p_ij_, is given by:

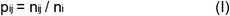

where n_ij_ is the number of transitions from note i to j, and n_i_ is the number of transitions out of note i. The first note in each sequence was chosen based on the relative frequencies of all notes in the dataset. Subsequent notes were chosen based on the transition probabilities from the previous note, as computed in the matrix M_0_.
3. We generated a new dataset from our original, where repeats of notes (i.e. self-transitions) were disregarded, thus each note could only transition to another note. Next, we generated the transition probability matrix for this dataset, denoted by M_1_, where all values along the diagonal were zero. For each note, we then fit the repeat distribution (the frequency of a note repeating once, twice, etc. normalized to the sum of observed frequencies) to a geometric distribution, where the probability of observing k repeats of any note, P_r_(k) is given by

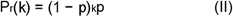

The parameter p was then fit separately to the repeat distribution for each note for each species. Thus, each note followed a unique repeat distribution in the model. Sequences were generated using the following algorithm: first, a note was chosen based on the relative frequencies of all notes in the dataset. Next, the number of times that note repeated was sampled from the geometric distribution described in (II). This number includes the case where the note only occurred once. The next note in the sequence was chosen with a transition probability given by M_1_, and finally the new note would have a probability of repeating as described above. To summarize, only transitions *between* notes were governed by a first-order Markov process, and the note could hold and repeat in the same state with the probability described in (II). This broadly matches renewal processes described in literature on animal vocal sequences (17). In general renewal processes, transitions between notes follow a first-order Markov process, but the note repetition distributions are governed by a Poisson distribution. We chose to use a geometric distribution instead, because all the notes had distributions with peaks at 1.
4. Finally, we generated sequences using a Hidden Markov Model (HMM) (57). This model consists of two components: hidden states that follow a first-order Markov chain, and emitted states which, in this case, are represented by the notes. The current hidden state of the model determines the probability of emission of different notes. Sequence structure is thus determined by two factors: 1) transitions between hidden states, and 2) the probabilities of uttering each note in the different hidden states. For example, consider an HMM with 2 hidden states A and B that generates sequences consisting of 2 notes, x and y. In state A, emission probabilities of x and y are (0.5, 0.5) whereas in state B, probabilities are (0.9, 0.1). The hidden state can transition from A to B (or B to A) with a probability 0.5, and the probability of occurrence of note x or y depends on whether the HMM is in state A or B. We obtained the number of hidden states for our model by minimizing the Akaike Information Criterion (AIC). We considered the number of states as the number of parameters while calculating AIC values. This was done in accordance with literature (17). Lower AIC values indicate a model that has both a good fit and low number of parameters (to avoid overfitting). We varied the number of hidden states from 2 to half the number of distinct note categories, and the AIC was calculated for each case. The number of hidden states that produced the lowest AIC value was selected to generate sequences.

Following this, we tested how well each model fit to the observed sequences using four primary methods. In the first method, we computed repeat distributions of each note for each species. A repeat distribution is the distribution of the proportion of occurrence of each repeat length (i.e. a note occurring singly, being repeated twice, thrice and so on) in vocal sequences before a transition to another note occurs, or the sequence terminates (5). We calculated the sum of squared errors (SSE) for each simulated repeat distribution against the observed distribution. The lower the SSE value, the better the fit of the simulated distribution (and thus the underlying process) to the observed distribution. This process was repeated for each note type, for each of the six species.

Next, we measured pairwise local alignment (the Smith-Waterman algorithm) of simulated sequences for each process against observed sequences (61). We assigned a score of +1 to matches and -1 to (i) mismatches, (ii) opening a gap, and (iii) extending a gap during alignment. Each alignment was associated with a score based on these penalties, and the optimal alignment was that with the largest (most positive) score. Larger alignment scores indicated higher sequence similarity, and thus a better match between simulated and observed sequences. To compare various models, we normalized the scores to account for larger sequences having higher scores on account of much greater lengths. This was done by dividing each pairwise alignment score by the natural logarithm of the sum of the lengths of the two sequences, to account for the great length variability in parrot vocal sequences. This method therefore works better to identify patterns in sequences of highly variable length, as opposed to other sequence similarity metrics that might show high dissimilarity simply on account of length variability.

Third, we used N-grams analysis, commonly used in animal communication and natural language processing, which examines frequency of occurrence of groups of N notes in a dataset (57,62). This analysis also rests on Markov assumptions, where an N-gram distribution simulates an (N-1)^th^ order Markov chain, and provides information on the occurrences of motifs or repetitive short structures within a sequence. The N-gram distributions were represented as multidimensional vectors, with each unique N-gram being one dimension. The component of the distribution vector along each dimension was the frequency of the corresponding N-gram. We then computed the Euclidean distance between each model’s distribution and that of the observed dataset for each species, thus enabling us to measure similarity. We performed this analysis for 2 and 5-grams. For each species, the model (and therefore, process) whose distribution had the lowest Euclidean distance from that of the observed dataset was considered the best fit.

Finally, we used the co-occurrence matrices defined previously as a metric to compare our models with the observed sequences. The co-occurrence matrices were generated with d=3, d=5 and d=7. For each case, we computed the Frobenius norm of the difference of the co-occurrence matrix for each model and the observed dataset, for each species. The Frobenius norm for two matrices *A* and *B* is given by

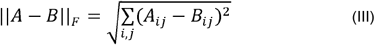

Essentially, this gave us the Euclidean distance between the two matrices, and a lower value implied a better fit of the model to the dataset.

## Results

### Note repetition in parrot vocal sequences

The number of note types emitted in sequences ranged from 5 for the two *Agapornis* species to 8 for

*P. krameri* (not including soft vocalizations ‘s’ that could not be classified to note type but were very rare in the dataset) (Figure 1). In all six species, co-occurrence values greater than chance typically existed along the diagonal of the note co-occurrence matrix, consistent with an elevated tendency to self-transition or repeat within a sequence (Figure 2).

**Figure 1:**
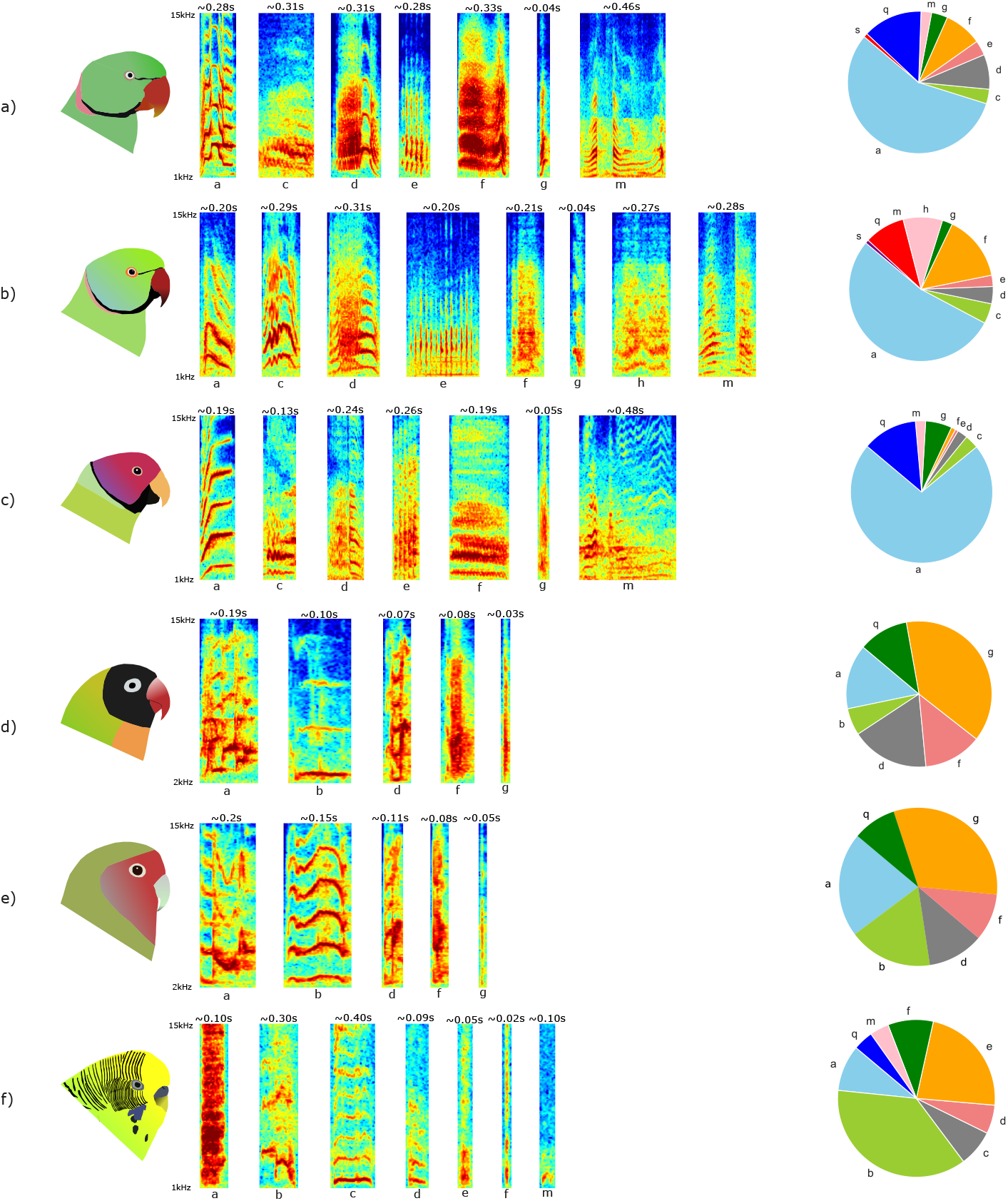
Vocal repertoires of six species within the Psittaculidae, with representative spectrograms showing the various note types, and the proportions of different note types in the vocal sequences of each species: a) Alexandrine Parakeet (*Psittacula eupatria*), b) Rose-ringed Parakeet (*Psittacula krameri*), c) Plum-headed Parakeet (*Psittacula cyanocephala*), d) Black-cheeked Lovebird (*Agapornis nigrigenis*), e) Rosy-faced Lovebird (*Agapornis roseicollis*), f) Budgerigar (*Melopsittacus undulatus*).

**Figure 2:**
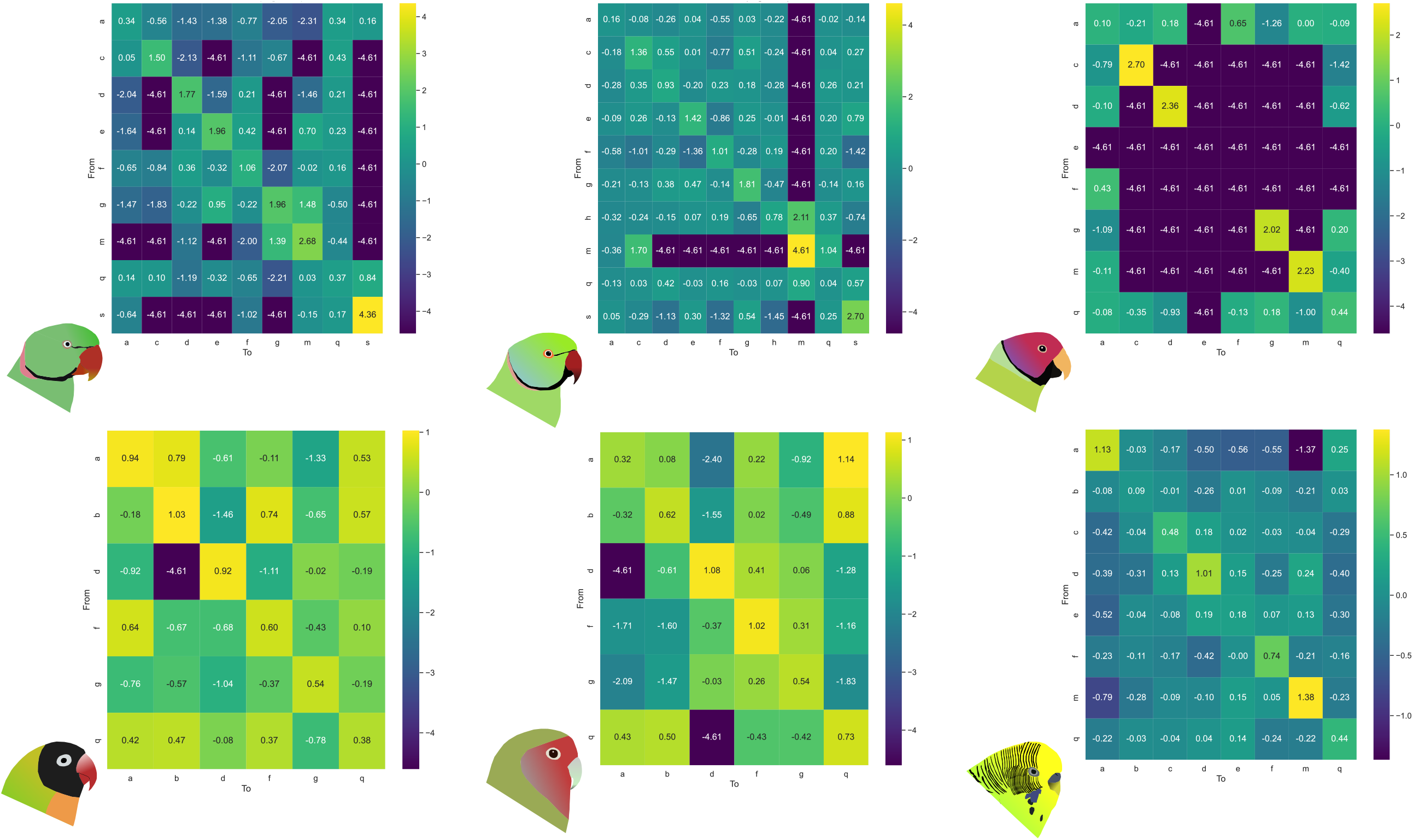
The matrices show the logarithm of observed:expected co-occurrence ratios of notes. A value greater than zero (warmer colors) implies the two notes co-occur more often than expected by chance, whereas one less than zero implies that the notes co-occur less often than expected by chance.

### Repeat distributions and local alignments are best explained by a renewal process

Next, we aimed to statistically test which process best characterized parrot vocal sequences, using diverse metrics (see Methods). First, we measured the SSE of repeat distributions (Figure 3) for each note type for each species. Broadly, for all six species, the repeat distributions for most notes fit best to the renewal process, and least to a random process (Figure 4). Although some notes for some species exhibited better fits to other processes, the broad trend across species was that the renewal process fit best to the maximum number of notes, and thus best explained the repeat distributions.

**Figure 3:**
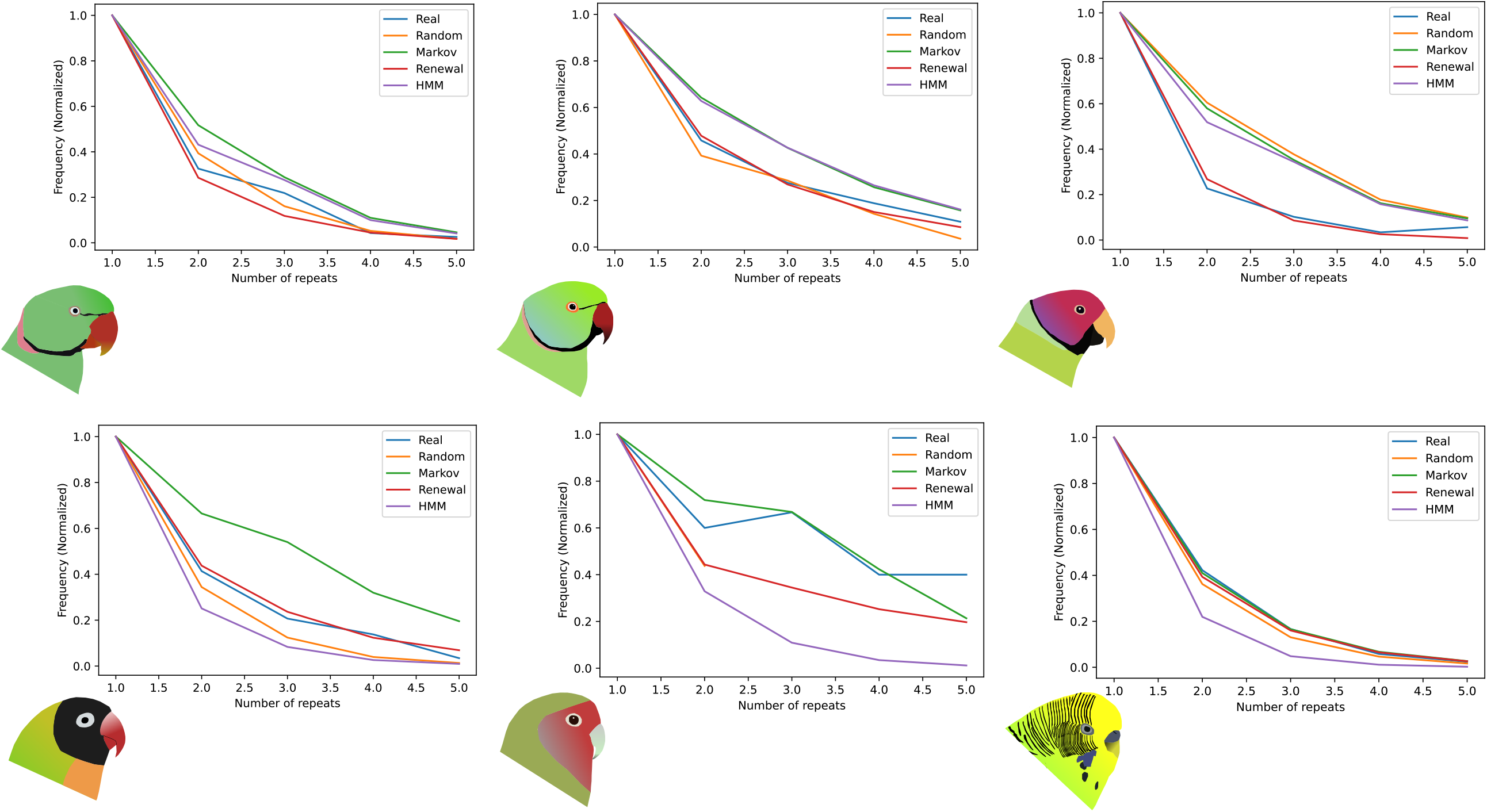
Example repeat distributions for the most common note type in each species, demonstrating the patterns obtained from observed sequences, as well as those expected under different simulated processes. The distribution shown for each species is that of the most abundant note in the repertoire: ‘a’ for *P. eupatria, P. krameri*, and *P. cyanocephala*, ‘g’ for *A. nigrigenis* and *A. roseicollis*, and ‘b’ for *M. undulatus*.

**Figure 4:**
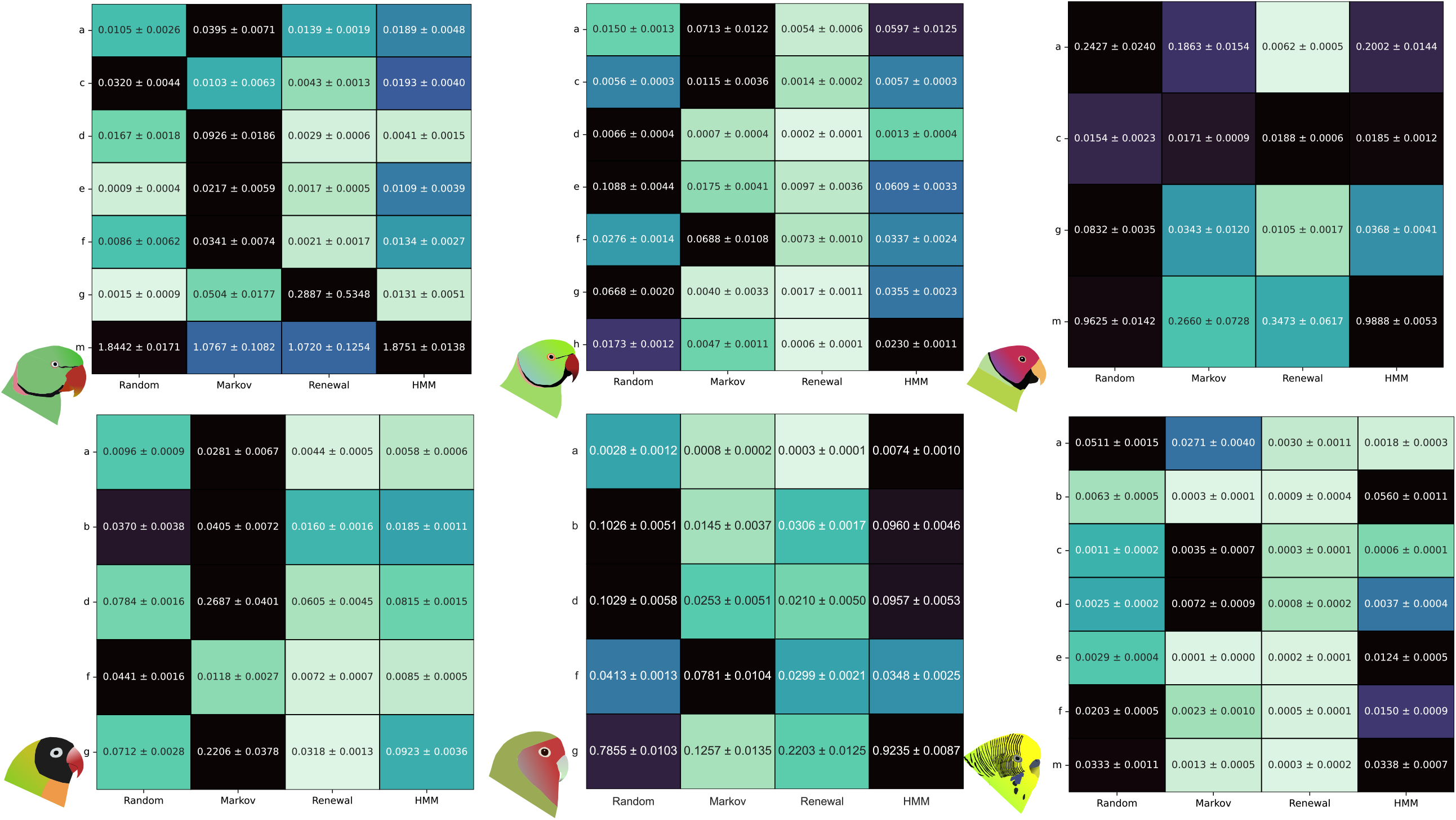
Summary of the results obtained by comparing observed repeat distributions to those obtained from four different simulated processes. Each figure shows the sum of squared errors (SSE) between the repeat distributions obtained from observed data and those from each simulated dataset. The lower the SSE (lighter colors), the better the fit.

We also used the strength of local alignments between observed and simulated sequences across species and different simulated processes. The renewal process fit best for 4 out of 6 species, whereas in the other two (*A. nigrigenis* and *P. cyanocephala*) the renewal and first-order Markov processes exhibited nearly identical local alignment scores (Figure 5).

**Figure 5:**
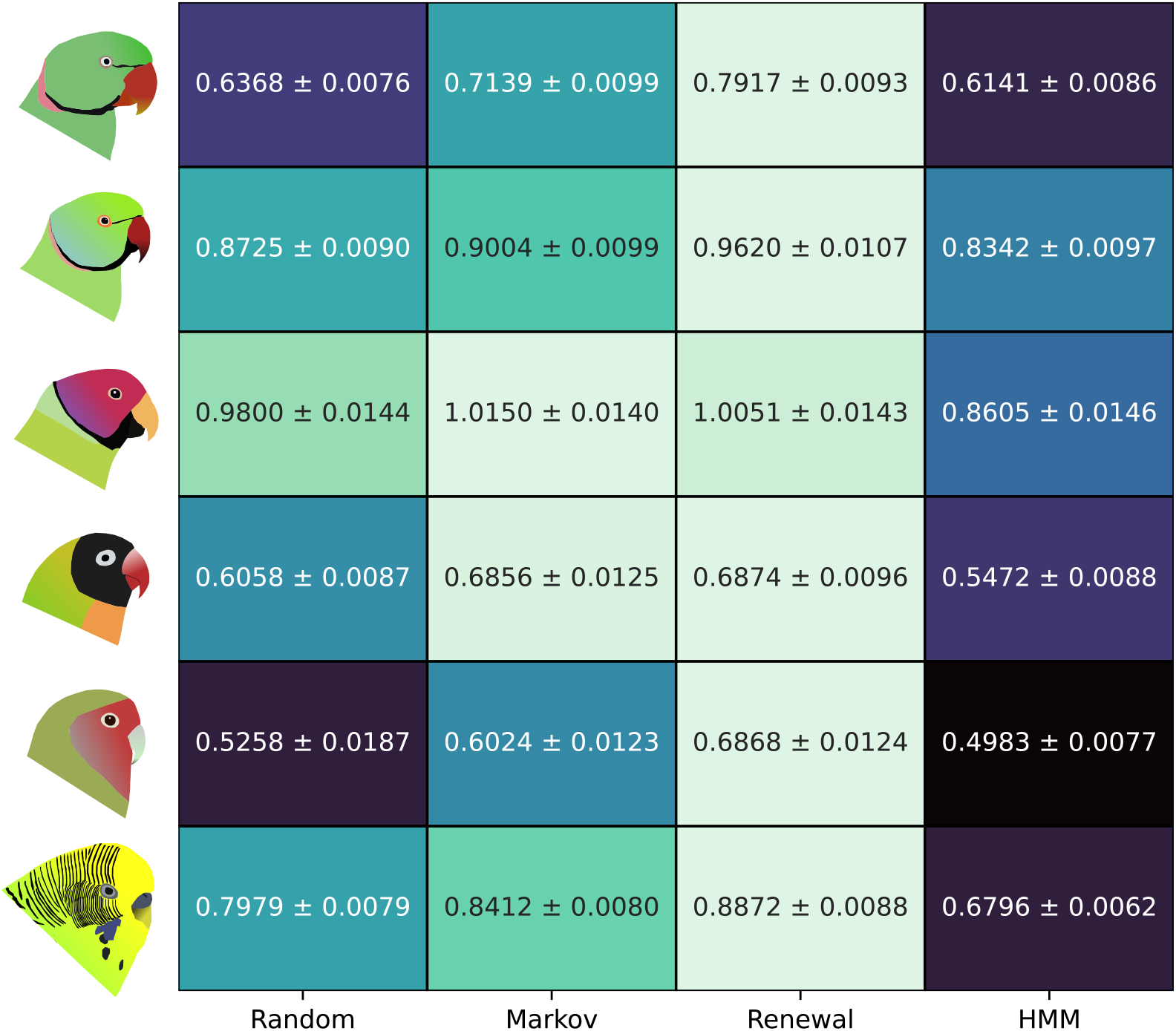
Local alignment scores between simulated and observed datasets, normalized to the lengths of the sequences (see Methods) for each species. The higher the score (lighter colors), the better the fit of that model to the dataset.

### Co-occurrence matrices and N-gram distributions favor first-order Markov processes

Next, we compared the observed co-occurrence matrices to simulated matrices generated across models by calculating the Euclidean distance between them (Supplementary Table 2). A lower value implied a closer match, and in all species the first-order Markov process best matched the co-occurrence matrix of the observed dataset, followed by the renewal process. This trend was observed for d=3, d=5 and d=7. However, the difference between the values for the two processes decreased with an increasing value of d (Figure 6), suggesting that considering smaller subsets of a sequence increased the tendency to favor a Markov process, with the apparent difference decreasing as the d-value was increased.

**Figure 6:**
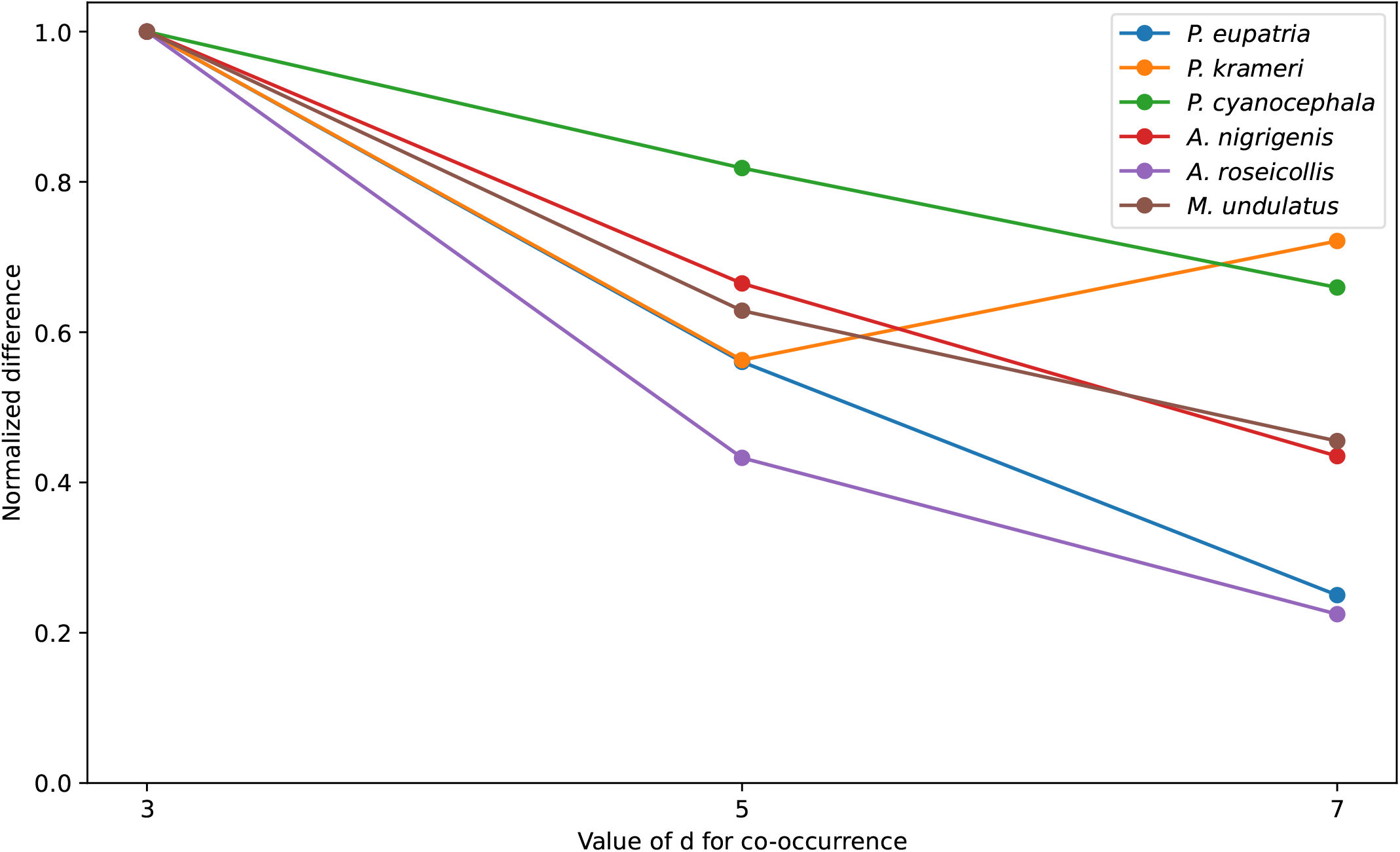
The Frobenius norm of the distance between two co-occurrence matrices. This distance was computed between each model and the observed dataset for each species. The normalized difference of this distance for the first-order Markov process and the renewal process are plotted here, for different values of d (see Methods).

Finally, we computed the distributions of occurrence of 2 and 5-grams for each model for each species and calculated the Euclidean distance between the model distributions and the observed data (Supplementary Table 3). The N-gram distributions favored the first-order Markov model consistently for both 2 and 5-grams (except for 5-grams distributions for *P. cyanocephala*, where multiple models performed comparably), albeit with larger Euclidean distances between observed and expected for 5-grams.

## Discussion

### Reconciling the results of diverse analyses

Because of the complexity of vocal sequences, the methodology used to examine patterns is diverse, drawing on the analytical frameworks of computational linguistics (7). Kershenbaum and Garland (2015) used a number of methods to examine similarity and patterns in sequences, both binary (sequence similarity measures) and unary (57). The best performing metric used Levenshtein edit distances between sequences as a metric of similarity. However, the length variability of sequences in parrots (often described as “rambling”) (3,41,63) means that this measure may inflate dissimilarities between observed and expected sequences. Therefore, while employing the same analytical rationale as the above study, we instead used local alignment, which provided measures of sequence similarity independent of the large variability in sequence length. In our study, diverse analytical methods uncovered similar patterns in the vocal sequences of different parrot species. Certain analyses supported the renewal process across species, whereas others supported a classical first-order Markov process. Importantly, we note that where one of these two processes was supported, the other usually performed second-best. Reconciling these results across species is somewhat complex, but similar patterns broadly hold true across all six species. This, at the very least, points to a general mechanism underlying vocal sequencing in parrots, one that places some emphasis on the role of note repetition (Figure 7). This last point is further buttressed by the co-occurrence matrices we constructed, which demonstrated an elevated tendency for notes to repeat themselves across all species (Figure 2). The analyses we used to examine patterns are widely used in linguistics. Why should different methods favor one process over another?

**Figure 7:**
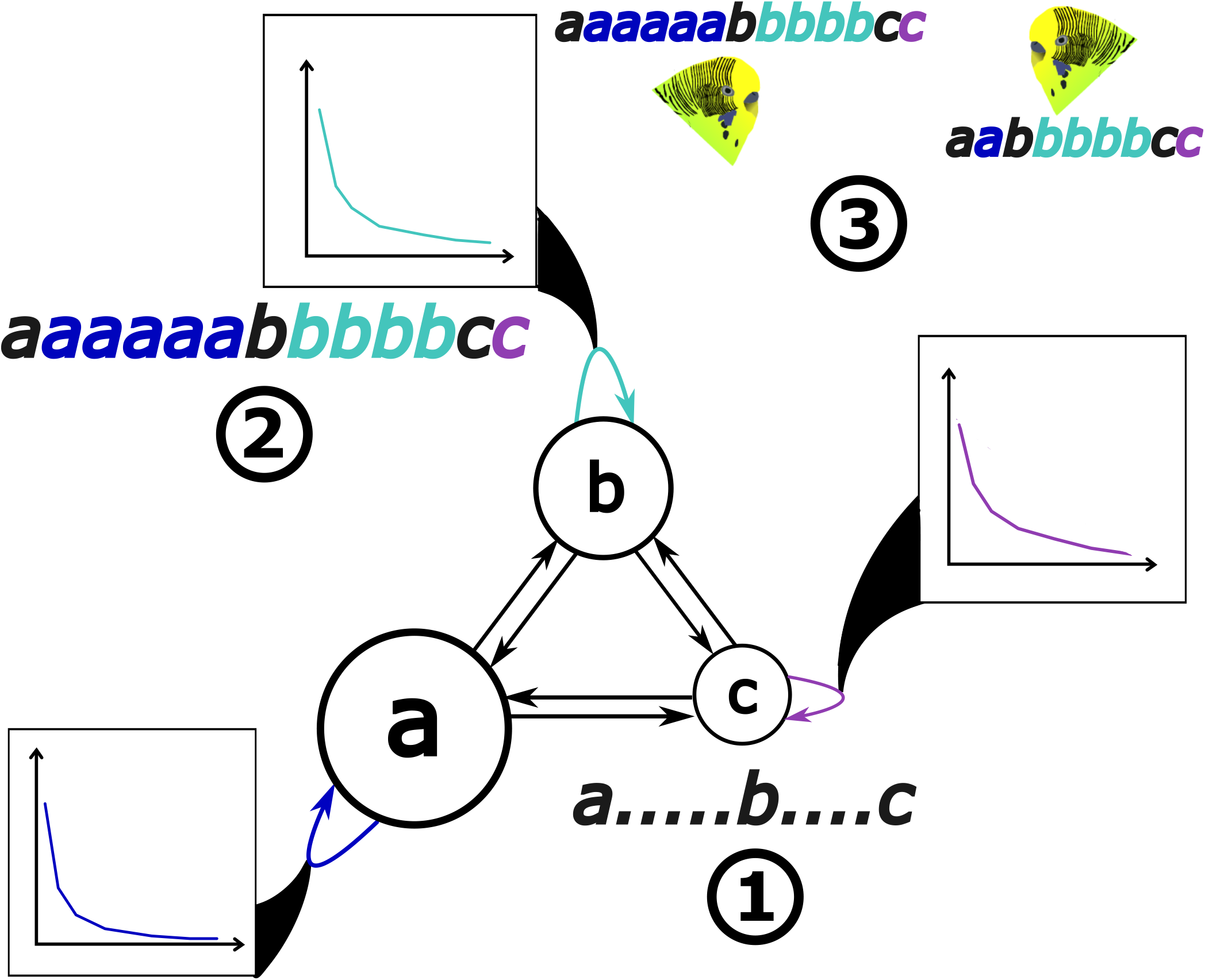
A schematic overview of note repetitions in budgerigars. **(1)** Transitions between different notes are governed by a Markov process. **(2)** A geometric distribution of repeat probabilities dictates how often each note repeats before moving to another note, as modeled by the renewal process. **(3)** Differing repeat distributions of specific notes may explain the discordance between analytical methods.

We suggest that the answer may arise from the underlying mathematical assumptions used to model each process. For example, N-grams analyses and co-occurrence matrices rely on capturing relatively short-range information depending on the size of N and the value of d used to generate the matrices (9,21,57,62), and this could explain why they tended to favor Markov processes. However, analyses using the local alignment scores potentially capture patterns over larger scales and tend to recover renewal as the best-performing process. Alternatively, patterns within sequences themselves may be responsible for the discordance across methods. In support of this, each metric broadly favored the same processes (e.g. local alignment scores favored renewal) across species. If note repetition alone is responsible for the variability of vocal sequences, then less repetitive sequences should perform similar to a first-order Markov process. The variation in repetitiveness and sequence length could explain why these two processes frequently perform similarly (or second to one another) in most of our analyses.

### The role of note repetition in conveying social information

Across species, co-occurrence matrices indicated an elevated tendency of notes to repeat themselves, with higher values along the diagonal (20). Both repeat distributions and local alignment scores indicated that renewal processes performed well, and thus supported the idea that notes in parrot vocal sequences tend to repeat themselves. Animal communication often exhibits redundancy, wherein multiple components of the same signal increase the probability of a receiver response (1). Repeating a note may similarly increase the probability of conveying information successfully. However, this hinges on the idea that each note is meaningful in and of itself (2,64,65), and that repetition would therefore be contingent on the location and responses of the receiver. The vocal sequences of budgerigars, at least, are long and very variable, and typically emitted during courtship to a nearby female recipient, where they may be associated with display movements (3,5,40–43). It is possible that female responses to repetitions of certain display steps may modify the occurrence of repeats in the song as a by-product (43).

The highly sociable, gregarious nature of most parrots (including all species studied here) means that vocalization also serves critical social functions (5,30,31,34–37,39,43,51,55). Multiple studies have shown that contact call structure may serve as a signal of group identity, rapidly changing by social learning when groups come into contact (31,34,35,48,49,53). Similarly, repeats of certain note types may serve as a syntactic signature of group identity, and these higher-order patterns also converge rapidly after groups come into contact (5). In this case, the repetition of notes may be specific across groups, and the presence of repeats may signal identity to members of the same flock. Further studies are needed on the social context of note repetitions, and on how different contexts modify the occurrence of repeats for different notes (7,36). Additionally, our analyses treated all notes within a repeat sequence as identical. However, contact calls and other notes exhibit within-category variability, and it is possible that spectral variability within repeat sequences encodes different types of information (66). Repetitions may thus increase sequence complexity within a limited repertoire, as variation within a note class may ensure that no two repetitions are similar. However, this requires further study, as does the perceptual role of these differences in repetition (40,41). Recent research demonstrates that budgerigars can discriminate differences in sequential structure of warble songs (63), which is consistent with findings of syntactic differences between groups (5).

## Conclusions

Open-ended vocal learners such as parrots provide a fascinating, albeit analytically challenging system in which to study vocal sequences (24). The presence of social learning together with complex fission-fusion flock dynamics introduces rapid sequence change (5,34,35,37,47), and the amenability of many parrot species for captive study (for example, in zoological collections) renders them uniquely suitable to examine the drivers and dynamics of social vocal change (37). Several of our analytical and computational methods, along with previous experimental evidence (5), highlight the important role that note repetition plays in parrots. As discussed above, note repetitions may serve many critical social functions, and we suggest that the combination of variability and repetitiveness in parrot vocal sequences may provide distinct results when tested against different processes. However, all species exhibited similar trends across analyses. This suggests that despite their complexity, there are some general organizing principles in parrot vocal sequences, in particular the importance of note repetition which was recovered using multiple analyses. By uncovering a general pattern across diverse members of the Psittaculidae from three different geographic regions (Africa, Asia and Australia) and different subfamilies (56), we highlight the potential for comparative insight to be gleaned from studies in zoological collections. We also highlight the need for a consensus among different analytical approaches when approaching a complex problem such as vocal sequence structure, as have other studies (57). The role of meaning and context in animal vocal sequences has been the subject of much discussion in the literature (7,17,57), and parrots present a tractable laboratory study system to test these hypotheses experimentally.

## Acknowledgments and funding

We are grateful to Dr. Uttam Yadav, Gaurav Patil, Shweta Pathak, Sangeeta Solanki, Aarti Xalxo and Ramesh Sawner for permissions and assistance during recordings, and the Galande family in Ujjain for accommodation and logistical support. All recordings of budgerigars were approved by the Institutional Animal Ethics Committee (Clearance AK 001) under the guidelines of the CPCSEA, New Delhi. AK is funded by a grant from the ANRF (CRG/2022/000187), Government of India, and AMD, NW, SS received fellowships from the KVPY program and NSD from the INSPIRE-SHE program, Government of India that supported research internships. The authors declare no competing interests.

## Supplementary Information

**Supplementary Table 1:** A summary of the note classification scheme for the species studied. The note classification for *M. undulatus* can be found in Madabhushi et al (2023) (5). Notes were considered distinct if they were separated by a gap of >10ms, and were thus distinctly separable from each other.

| Species | Note type | Description |
| --- | --- | --- |
| *P. eupatria*, *P. krameri*, *P. cyanocephala* | a | Contact calls and harmonics |
|  | c | Wavy calls |
|  | d | Harsh harmonics (>25% of the duration of the note is harsh) |
|  | e | Trills (a rapid sequence of short clicks) |
|  | f | Harsh (broadband non-harmonic calls) |
|  | g | Clicks (broadband calls shorter than 60ms) |
|  | h (*krameri* only) | Gruff/throaty harmonics |
|  | m | Mimicry (cases where the parrot uttered human words) |
|  | s | Soft calls (too low in amplitude to classify) |
| *A. nigrigenis*, *A. roseicollis* | a | Chirp-like (multiple jumps in frequency) |
|  | **b** | Pure harmonics |
|  | **d** | Harsh harmonics (any duration of the note, however small, is harsh) |
|  | **f** | Harsh (broadband non-harmonic) |
|  | **g** | Clicks (shorter than 55ms for *nigrigenis* and 65ms for *roseicollis*) |

**Supplementary Table 2:** The Euclidean distances between co-occurrence matrices for the observed dataset and each model are shown here for each species (see Methods). Across species, the first-order Markov model performs best, with the renewal process performing second best. However, with increasing values of d, the difference between the first-order Markov and the renewal process decreases.

|  | **d=3** | **d=5** | **d=7** |
| --- | --- | --- | --- |
| ***P. eupatria*** | Random: 62.8091 ± 3.1419<br>Markov: 24.2952 ± 1.0507<br>Renewal: 53.8264 ± 3.0366<br>HMM: 63.4432 ± 3.3300 | Random: 50.1477 ± 1.8124<br>Markov: 27.8748 ± 0.7781<br>Renewal: 44.4250 ± 1.6579<br>HMM: 51.6873 ± 1.6823 | Random: 42.0550 ± 0.7958<br>Markov: 28.1633 ± 0.6147<br>Renewal: 35.5488 ± 0.6999<br>HMM: 40.3467 ± 0.9241 |
| ***P. krameri*** | Random: 5.0261 ± 0.0139<br>Markov: 1.7812 ± 0.0031<br>Renewal: 2.5648 ± 0.0035<br>HMM: 6.6667 ± 0.0099 | Random: 4.5121 ± 0.0060<br>Markov: 2.6488 ± 0.0036<br>Renewal: 3.1698 ± 0.0028<br>HMM: 6.6065 ± 0.0161 | Random: 4.5602 ± 0.0064<br>Markov: 3.1909 ± 0.0031<br>Renewal: 3.5318 ± 0.0029<br>HMM: 6.5172 ± 0.0135 |
| ***P. cyanocephala*** | Random: 5.2420 ± 0.0218<br>Markov: 2.0873 ± 0.0112<br>Renewal: 2.9080 ± 0.0109<br>HMM: 8.3540 ± 0.0212 | Random: 4.1250 ± 0.0182<br>Markov: 2.2370 ± 0.0075<br>Renewal: 2.7529 ± 0.0079<br>HMM: 7.9488 ± 0.0142 | Random: 3.5311 ± 0.0121<br>Markov: 2.1974 ± 0.0064<br>Renewal: 2.5709 ± 0.0075<br>HMM: 7.7994 ± 0.0139 |
| ***A. nigrigenis*** | Random: 18.9500 ± 0.0982<br>Markov: 10.3988 ± 0.0768<br>Renewal: 15.8155 ± 0.1067<br>HMM: 33.0117 ± 0.4944 | Random: 19.9370 ± 0.1091<br>Markov: 12.9192 ± 0.0587<br>Renewal: 17.3522 ± 0.0702<br>HMM: 32.1810 ± 0.3223 | Random: 19.2514 ± 0.0734<br>Markov: 13.9583 ± 0.0577<br>Renewal: 17.5304 ± 0.0918<br>HMM: 31.3043 ± 0.1996 |
| ***A. roseicollis*** | Random: 30.1918 ± 1.7342<br>Markov: 13.9064 ± 1.1907<br>Renewal: 17.1553 ± 3.1640<br>HMM: 436.9011 ± 39.8476 | Random: 24.6887 ± 2.1995<br>Markov: 14.2217 ± 1.0511<br>Renewal: 16.0503 ± 3.5518<br>HMM: 431.2610 ± 30.5632 | Random: 21.4424 ± 2.4145<br>Markov: 11.1624 ± 0.4639<br>Renewal: 13.5060 ± 3.8589<br>HMM: 414.3002 ± 30.6009 |
| ***M. undulatus*** | Random: 8.0606 ± 0.0094<br>Markov: 1.7839 ± 0.0016<br>Renewal: 2.4531 ± 0.0057<br>HMM: 6.6559 ± 0.0097 | Random: 7.4578 ± 0.0065<br>Markov: 2.3637 ± 0.0033<br>Renewal: 2.6533 ± 0.0028<br>HMM: 6.1003 ± 0.0059 | Random: 7.0170 ± 0.0049<br>Markov: 2.6973 ± 0.0025<br>Renewal: 2.8477 ± 0.0022<br>HMM: 6.1519 ± 0.0063 |

**Supplementary Table 3:**
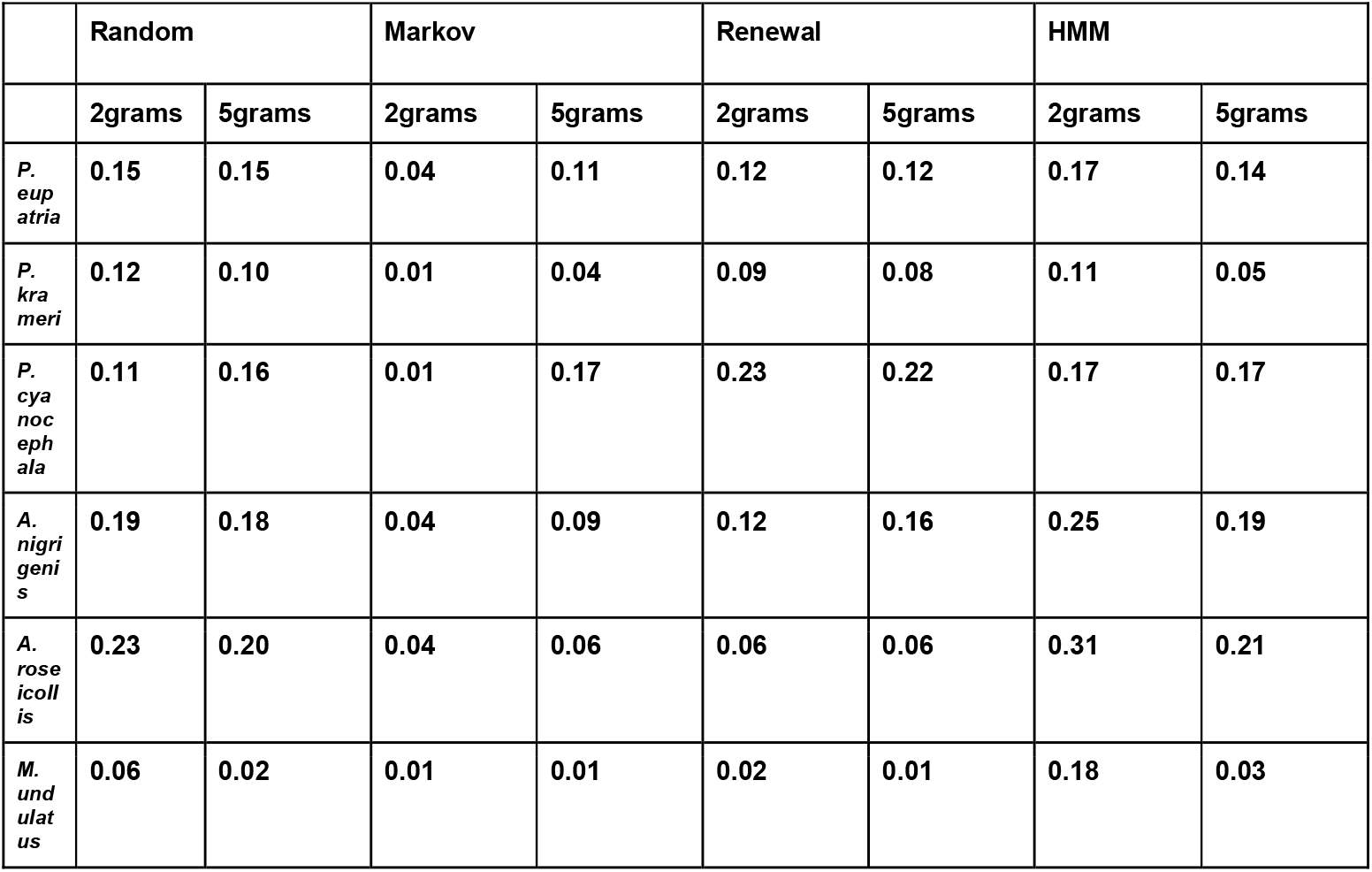
The Euclidean distances between N-gram distributions for the observed dataset and each model are shown here for each species (for N = 2 and 5) (see Methods). Across species, the Markov model is the closest to the observed dataset (lower Euclidean distance). For 5-grams, the Markov model still performs best for all species (excluding *P. cyanocephala*), but the Euclidean distances are larger than for 2-grams.

